# Cracking the problem of neural representations of abstract words: grounding word meanings in language itself

**DOI:** 10.1101/391052

**Authors:** Annika Hultén, Marijn van Vliet, Lotta Lammi, Sasa Kivisaari, Tiina Lindh-Knuutila, Ali Faisal, Riitta Salmelin

## Abstract

In order to describe how humans represent meaning in the brain, one must be able to account for not just concrete words but, critically, also abstract words which lack a physical referent. Hebbian formalism and optimization are basic principles of brain function, and they provide an appealing approach for modeling word meanings based on word co-occurrences. Here, we built a model of the semantic space based on word statistics in a large text corpus, which was able to decode items from brain signals. In the model, word abstractness emerged from the statistical regularities of the language environment. This salient property of the model co-varied, at 280–420 ms after word presentation, with activity in the left-hemisphere frontal, anterior temporal and superior parietal cortex that have been linked with processing of abstract words. In light of these results, we propose that the neural encoding of word meanings is importantly grounded in language through statistical regularities.

## Introduction

Understanding abstract and concrete concepts is a fundamental aspect of human language that enables us to discuss matters ranging from everyday objects to fantastic stories of fiction. A common view is that word meanings are grounded in experiences with the world [1–4]. For example, the word ‘tomato’ is linked with the look, feel and taste of a tomato. This view of lexical semantics assert that these types of physical associations form the building blocks of how words are encoded in the brain. However, the grounding framework fails to account for abstract words, which lack physical referents and, in many cases an emotion or an internal state to which the word meaning can be grounded. This issue can be overcome, if word meanings can also be grounded in the experience of language. In this conceptualization, representations of both concrete and abstract words will mirror, not merely physical, but any regularities in the environment. We address this question starting from the framework of statistical self-organization of the semantic space and aim to demonstrate that both concrete and abstract words can be represented in the brain based on their statistical properties that emerge directly from natural language.

Computational models of in the field of natural language processing (NLP) have demonstrated that a distributed representation of word meanings can be derived from the context in which the words are used. The core idea of these models is to find an optimal decomposition of semantics that can represent each unique concept without excessive use of memory or processing effort. Statistical regularities in the training data (typically a large text corpus) will drive the organization of the semantic space, wherein categorical structures, such as that of abstract and concrete words, can emerge [5]. These models rely on the same general computational principles that underlie brain function, namely Hebbian learning [6] and basic principles of optimization [7, 8]. If we further assume that a large text corpus is a fair estimate of the natural language environment that our brains are immersed in, a statistical model of a text corpus could serve as a reasonable approximation of the organizational principles of word meanings also at the level of the brain. In this framework, statistical regularities in the language environment are mirrored in the semantic space of the brain, and neural representations of both abstract and concrete words are thus formed by the same general computational principles.

Systematic patterns in the language environment can give rise to qualitative differences in the way concrete and abstract words are represented or processed, even if those word types share the same organizational principles. Behaviorally, concrete words elicit faster reaction times than abstract words [9]. Patient data suggest a double dissociation between abstract and concrete word types as either one may be selectively impaired [10, 11]. Furthermore, numerous neuroimaging studies have shown that processing of abstract and concrete words activate brain areas differently [12]. Generally, processing of abstract words (nouns in particular) activates classical language areas, such as the inferior frontal gyrus and the middle/superior temporal gyrus, more strongly than processing of concrete words. In contrast, concrete words seem to activate the posterior cingulate, precuneus, fusiform gyrus, and parahippocampal gyrus more strongly than abstract words (ibid). Electrophysiological evidence reports stronger and longer-lasting neural response for concrete than abstract words at around 400 ms after word onset [13, 14].

Here we will show that it is possible to explain cortical activation to a wide selection of abstract and concrete words using a model derived only from the statistical relationships among words in our natural language, without summoning qualitative differences in the acquisition of the words. A conceptually straightforward and computationally explicit model is used to describe emergent categorical organization and the way in which the meaning of words can be represented in the brain. We focus on the emergent properties that arise through self-organization (self-organizing map [15], SOM) in an artificial neural network model of the language environment [16], and use magnetoencephalography (MEG) to examine whether this structure is mirrored in brain activity during word reading.

## Results

The Statistical model of word meanings was built by applying the Word2vec algorithm to a large text corpus of the Finnish internet. The algorithm was developed in the field of natural language processing [16, 17], and it bases its notion of semantic similarity on the principle that two words are similar if they occur within a similar linguistic context, even if they never directly co-occur. Word2vec will discover thematic relationships (bear – zoo), i.e., concepts that either serve complementary roles or that co-occur in common situations, locations and/or times, but do not necessarily share perceptual or functional characteristics [18, 19].

In order to assess the internal structure and emergent properties of the Statistical model, we visualized the relationships among 118 MEG stimulus words (59 abstract and 59 concrete nouns) using an unsupervised SOM [15, 20]. This method provides a two-dimensional representation of the distances from each word to all other words in the semantic model.

The SOM visualization of the corpus-derived Statistical model (see Figure 1) revealed a clear division between abstract and concrete words. The abstract words (e.g., freedom, ideology, democracy) group together and are distinct from the concrete words (e.g., castle, tower, bridge). Notably, the words rated as medium abstract (e.g., music, song, poem) indeed appear to be situated between the concrete and highly abstract words in the Statistical model. The map also shows that the organizational principle of the concrete words does not strictly align with the six a priori defined taxonomic categories. For example, the words in the Human character category (e.g., police, prisoner) are interleaved with words classified as belonging to Building (e.g., prison) in a manner that seems to capture a thematic association. Only the taxonomic categories Body parts and Objects appeared as clearly distinct groups in the Statistical model.

**Figure 1:**
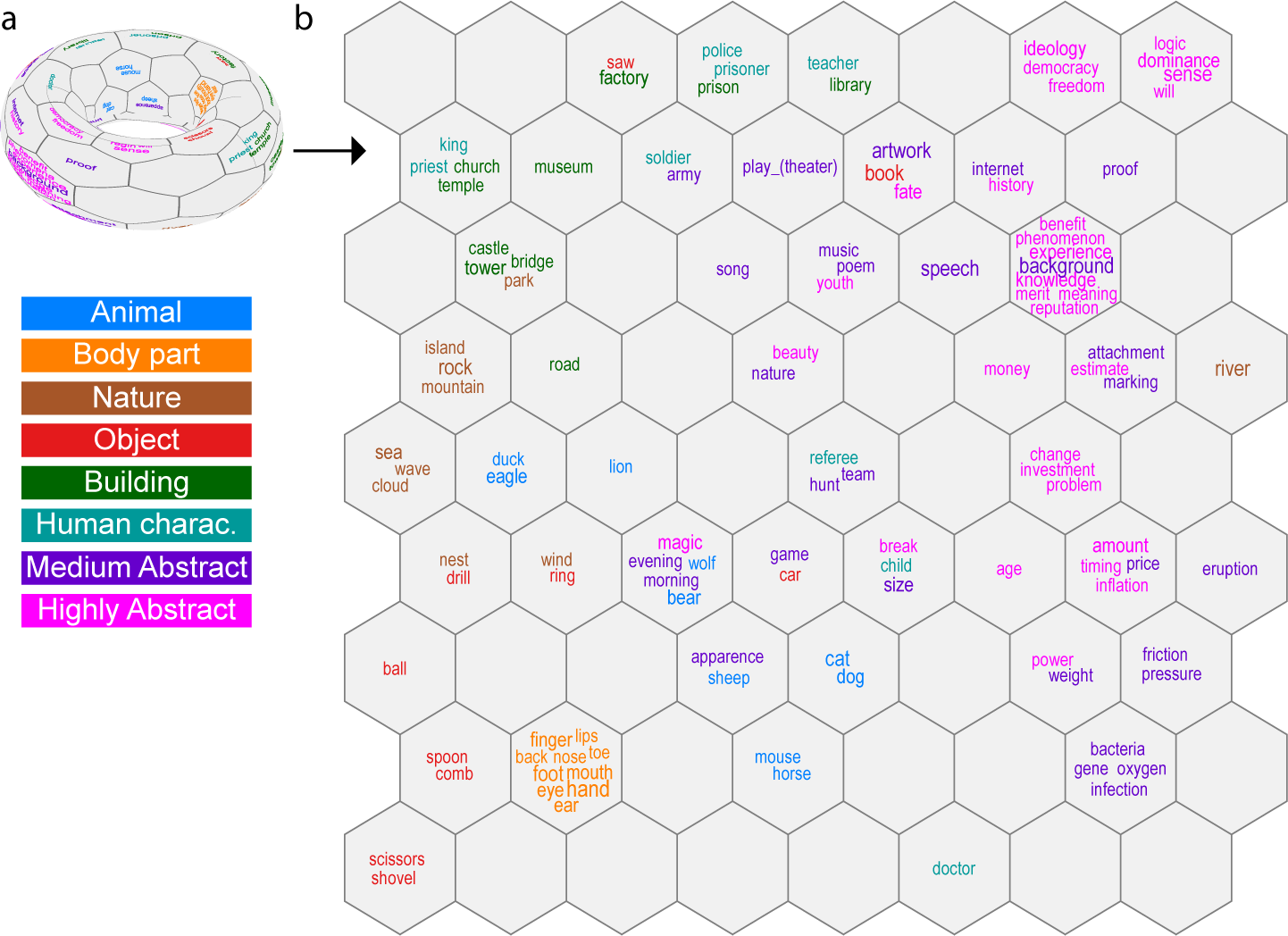
Self-organizing map of the Statistical model. The map is modeled on the surface of a toroid (A), with a continuous network of the nodes. For visualization, the toroid is “cut” and unfolded into a two-dimensional sheet (B). The size of the each word denotes the distance to the node center (the larger the font, the closer to the node center the word lies) and the color the predefined class of the stimulus words. The grouping of the words in a SOM should be interpreted based on the size of the clusters and the relationship between a particular grouping and other groups. The exact position of a word on a SOM visualization is generally not informative.

To determine whether the information in the Statistical model is mirrored in the brain activity during word reading, the activation evoked by each word was measured with MEG from 20 volunteer participants. The participants read the words, which were presented one at the time in a random order and repeated 20 times each over the course of the experiment. Evoked responses were formed by averaging the neural response across the 20 presentations of each stimulus word, for epochs starting −200 ms before word onset until 1000 ms after it.

An item-level decoding algorithm [21] was used to evaluate how well the corpus-derived Statistical model serves as a model of brain-level organization of word representations. This supervised machine-learning model tries to find an optimal linear mapping between the MEG data (204 sensors × 40 time bins) for each stimulus word and the corresponding feature decomposition of the word from the Statistical model [22]. The success of the machine-learning model is evaluated using a leave-two-out cross-validation scheme, i.e., asking the model to distinguish between two words that it had not previously encountered (116 words used to train the model, 2 words left out for testing, all permutations). The left-out words were successfully discriminated in the majority of participants (15/20 participants), with a mean prediction accuracy across all participants at 63.6 % (s.d. 7.5) and a top score of 77.8 % (Figure 2). The adjusted chance level was determined statistically to be 60.1 % (*p* < 0.05). The algorithm was thus able to find a mapping between the brain data and the Statistical model, which implies that the information encoded in the Statistical model is correlated with the information in the brain signal. A breakdown of the item pairs used in the evaluation showed no between-category advantage compared to within-category comparisons, indicating that the decoding accuracy was not merely driven by the categorical structure of the stimulus words (see Supplementary Figure S1).

**Figure 2:**
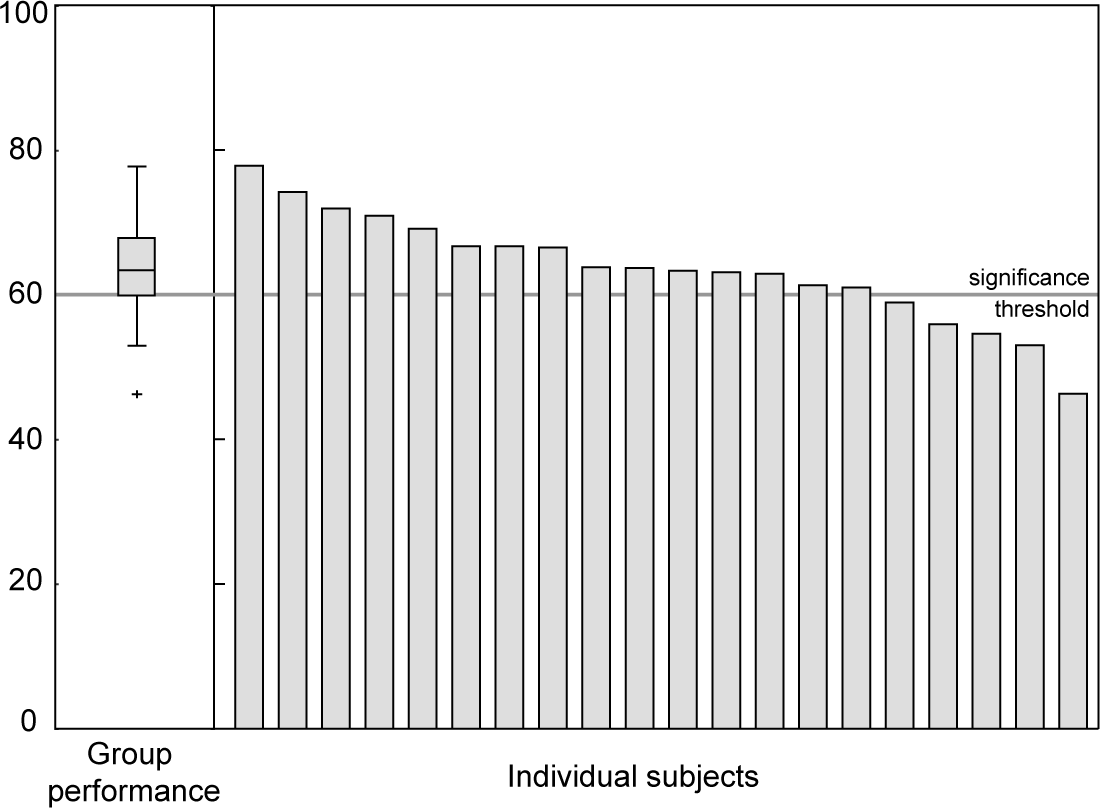
Item-level decoding accuracy. The box plot on the left shows the quartiles and the variation in the group performance (percent of successful decoding across all item-set permutations). On the right are the individual scores of each participant. The accuracy scores above 60 % were deemed statistically significantly above the chance level based on a permutation test.

We proceeded to investigate when and where the information expressed in the Statistical model is manifested in brain activation by using representational similarity analysis [23] (RSA) between the MEG data and the semantic decompositions provided by the Statistical model (Figure 3A). The RSA aims to discover time bins and cortical regions where the variation in the source estimate of the MEG signal is similar to the variation in the model. To determine associations that are consistent across the 20 participants, the significance of the RSA maps was evaluated using a cluster permutation test [24].

**Figure 3:**
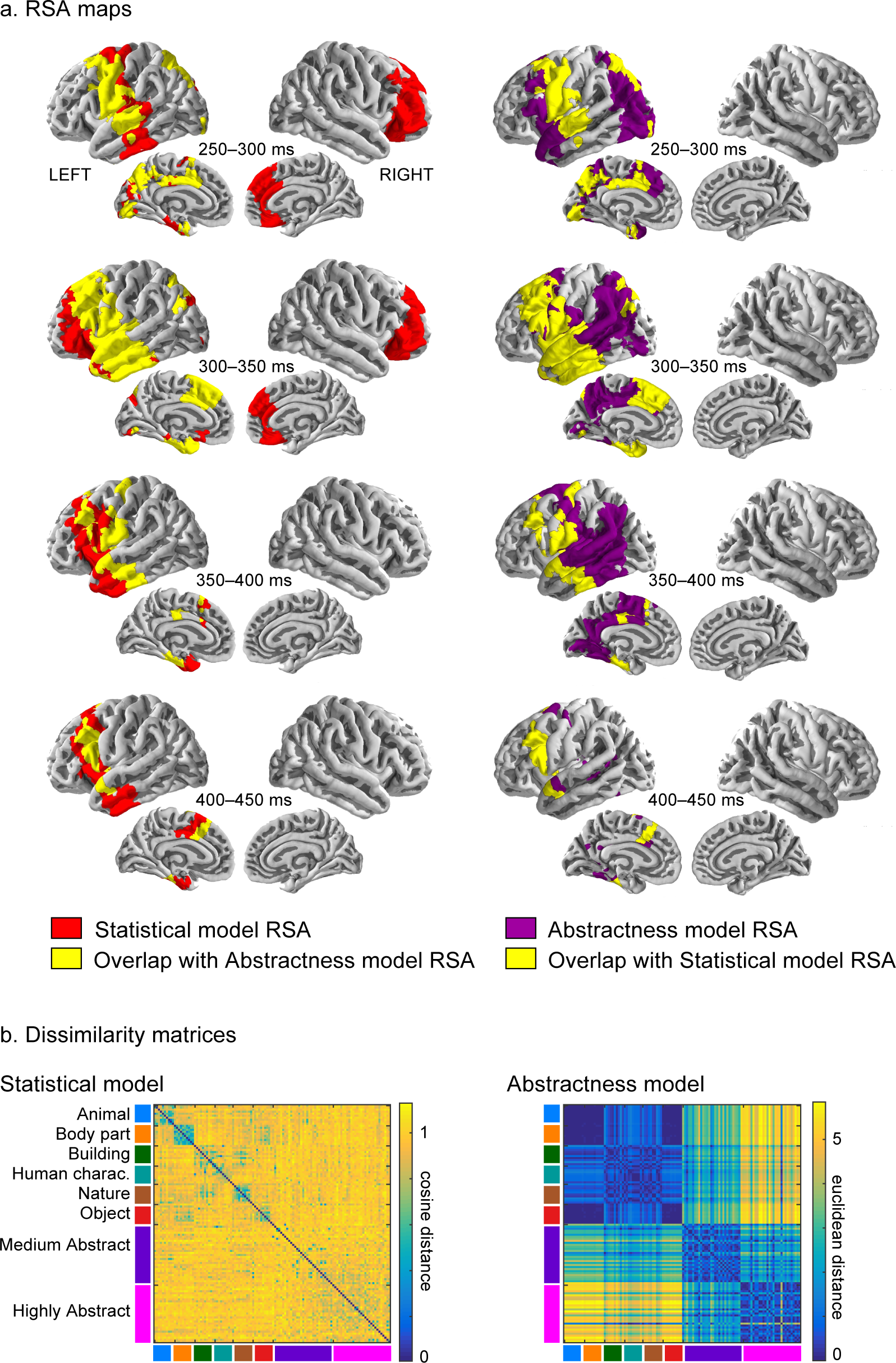
Comparison of the Statistical model and Abstractness model. (A) Representational similarity analysis (RSA) between the Statistical model and the MEG data (red) on the left and between the Abstratcness model and the MEG data (purple) on the right. The overlap between the two RSAs is plotted in yellow. The results show all regions and time windows with statistically significant findings. For visualization purposes, the data was averaged over 50 ms time windows. (B) Dissimilarity matrices of the Statistical model and the Abstractness model.

Based on the SOM analysis, the dominant organizational principle of the Statistical model is the abstractness-concreteness dimension. Therefore, to guide the interpretation of the RSA between the MEG data and the Statistical model (henceforth, Statistical model RSA), we additionally calculated a complementary RSA between the MEG data and a model used to quantify only the abstractness – concreteness dimension based on independently collected questionnaire data (see Methods; henceforth, Abstractness model RSA). The questionnaire data (Abstractness model) clearly dissociated between concrete, medium abstract and highly abstract words (see dissimilarity matrices in Figure 3B). The spatio-temporal overlap between the Statistical model RSA and Abstractness model RSA (Figure 3A) suggests that neural activity in the highlighted cortical regions contains information incorporated in both model types. A large portion of this information is related to the abstractness dimension, as shown by the high correlation between the Abstractness model and the Statistical model (Mantel test with Spearman’s *ρ* = 0.3, *p* < 0.001; 5000 permutations used). However, the RSA cannot distinguish if the strength of brain activity is larger at one end of the spectrum (e.g., whether abstract words elicit greater brain activation than concrete or vice versa).

The earliest neural response that was significantly correlated with the Statistical model was observed in the left precentral gyrus (280 ms to 340 ms), with activation subsequently extending across large parts of the frontal cortex, including the left inferior frontal gyrus (320 ms to 420 ms) and medial superior frontal cortex (320 ms to 360 ms). Activity patterns in the left temporal cortex were also significantly correlated with the Statistical model; first in the middle part of the superior temporal cortex (280 ms to 340 ms), followed by the medial (300 ms to 240 ms; 380 ms to 420 ms) and lateral parts (320 ms to 420 ms) of the anterior temporal lobe.

There was substantial overlap between the Statistical model RSA and the Abstractness model RSA starting from the inferior precentral gyrus (300 ms and superior middle temporal cortex. In a later time window, the overlap between the two RSA analyses in frontal lobe extended to the inferior, middle, superior and medial regions (320 ms to 400 ms). The Statistical model RSA additionally found correlation in the left precuneus and superior parietal cortex (280 ms to 340 ms) that partially overlapped with correlations in these areas found in the Abstractness model RSA.

The only areas highlighted uniquely by the Statistical model RSA throughout the timeline were found in the right frontal cortex (280 ms to 340 ms). Activity in the left posterior temporal cortex and temporo-parietal junction (280 ms to 380 ms) was only significantly correlated with the Abstractness model.

## Discussion

We showed that abstract and concrete words self-organize into distinct groups based on their occurrence in different sentential contexts. Notably, this categorical organization principle emerges through basic principles of association and optimization of the statistical information present in the language environment. The alignment of words rated as medium abstract between the concrete and abstract words in the semantic space suggests the abstract-concrete distinction is a continuum.

The statistical properties of word meanings derived though co-occurrences in the text corpus were successfully used to decode the identity of written words based on their MEG responses, thereby showing that a statistical model of semantics can be used to explain the organization of word meanings in the brain. The main correlations between brain activity and the Statistical model were found in the left precentral, frontal and anterior temporal cortex as well as in superior and medial parietal areas. We interpret the findings in light of the complementary model that expresses the level of abstractness, focusing on the brain areas revealed by both the Statistical model RSA and the Abstractness model RSA.

The overlap discovered between the Statistical model RSA and Abstractness model RSA is in line with the common finding that processing of abstract words (nouns in particular) activates classical language areas, such as the inferior frontal gyrus and the middle/superior temporal gyrus, more strongly than processing of concrete words (for a meta-analysis, see [12]). The left inferior frontal gyrus in particular, has been highlighted as an especially informative area in decoding the abstract/concrete word class [25]. In the present study, overlap between the two RSA maps was observed both in the inferior/superior left frontal cortex (320 ms to 420 ms) and in the superior temporal cortex (300 ms to 340 ms). In previous studies, more activity for abstract and concrete words in these areas was interpreted to reflect greater engagement of the verbal system for processing of abstract concepts [12]. It is therefore not surprising that the information processed in this classical language area mirrors both the statistically derived categorical structure and especially the abstract-concrete dimension. The present RSA findings further suggest that the adjacent superior and medial parts of the frontal cortex are also able to capture the abstractness dimension of word meanings.

The present RSA results also identified areas that in previous studies have shown stronger activation to concrete than abstract words [12] or displayed strong linkage to item-specific semantics of concrete words [26], namely the posterior cingulate, precuneus, fusiform gyrus, and parahippocampal gyrus. In previous studies, increased activation for concrete words has often been interpreted in terms of grounding conceptual information to the perceptual system, particularly in the ventral or dorsal visual processing streams [12, 27]. The present findings show that the patterns of activation in these areas is also correlated with the Statistical model derived from corpus data.

Furthermore, the Statistical model RSA and Abstractness model RSA both revealed semantic encoding in the lateral and medial parts of the anterior temporal lobe at 320 ms to 420 ms (including ventro-medial regions). The anterior temporal cortex is well-known for its role in both semantic dementia [28] and associative semantics [29]. In light of the associative nature of the statistical semantic model, the present results support the notion that this region is in some manner also linked with processing of word meanings through their associative properties to other words.

A prominent overlap between the Statistical model RSA and Abstractness model RSA was additionally observed in the left precentral cortex (300 ms to 340 ms). This region has previously been linked to category-specific semantic activation related to body parts and shape [30]. Here we show that the activity pattern in this region also aligns with the abstract-concrete structure in the Statistical model.

Self-organization of the semantic space provides an account for how differences along the abstract-concrete dimension can arise. Assuming that the neural representations of word meanings arise from similar computational principles as the ones that govern the Statistical model, words that co-occur in the environment will also share some aspects of their neural representation. This could lead to categorical groupings and thus explain the categorical differences that have been found in previous experimental and clinical studies [12].

Most of the cortical areas discovered in the RSA analyses align with classical language areas, outside of the primary motor or sensory areas. This suggests that the abstractness dimension is more than a mere reflection of direct sensory-motor associations, put forward by some advocates of the embodied cognition view [1, 2]. This finding would explain why previous attempts at decoding abstract words based on sensory-motor attributes have been unsuccessful [31] whereas even a crude nominal categorical classification of the abstractness dimension seems to work [25]. When using a more detailed description of the semantic space, such as the present corpus-derived Statistical model, we were able to decode MEG signals of individually presented written words; the written modality has previously proven to be challenging even in categorical classification of concrete words [32].

The choice of semantic model used to describe the semantic space matters. Despite the marked overlap between the Statistical model RSA and the Abstractness model RSA, several areas were uniquely highlighted by only one of the models. This suggests that the Statistical model does not capture all aspects of the abstract-concrete dimension (or these aspects may remain below the statistical significance threshold). Similarly, while the Word2vec model is a well-argued model of item-level semantics, alternative models such as those based on behavioral feature descriptions may provide complementary views to the semantic system.

The present study provides a computationally explicit framework of how semantic representations can be expressed in the brain, in the form of a statistical model that uses computational principles known to exist in the brain. We were able to link specific cortical areas to semantic representations but also to describe the type of information that could be processed there and how it may have come about. We show that a statistical model is sufficient to explain a substantial part (i.e., enough to enable successful encoding) of the semantic processing. This suggests that the experience of language can be seen as equivalent to any other sensory, motor, emotional or perceptual experience. Abstract words could, therefore, be grounded in language itself, making second order grounding through, e.g., metaphors redundant.

## Methods

### Participants

MEG measurements were performed on 20 volunteers (mean age 21 years, sd 3.6, range 18–34; 50 % identified themselves as females). All participants were native Finnish speakers, had normal or corrected to normal vision, and were scored as highly right-handed on the Edinburgh handedness questionnaire. The participants were all healthy, reported no diagnosed neurological disorders or reading disabilities and were compensated financially for their participation. Informed consent was obtained from all participants. In addition, a total of 408 respondents filled behavioral questionnaires, created either for stimulus evaluation or to collect the behavioral feature sets (see more information below). All the respondents were volunteers who were reimbursed for the effort with movie tickets. All respondents had Finnish as their first language, their mean age was 27 years (sd 7, range 19–63) and 65 % identified themselves as females.

The study was approved by the Aalto University Research Ethics Committee in agreement with the Declaration of Helsinki.

### Stimuli

The stimuli consisted of 118 nouns grouped into two main categories: concrete (59 words) and abstract (59 words). The two main categories did not differ significantly in lemma frequency [*t* (58) = −1.1, *p* = 0.28], based on the prevalence in Finnish-language internet pages (corpus size 1.5 billion words). All words were within the 90th percentile of the corpus distribution and can thus be considered common, high frequent words. The length of the stimulus words ranged from 3 to 10 letters and did not differ between the abstract and concrete words [*t* (58) = −1.9, *p* = 0.065].

All stimulus words were assessed on a scale from 1 to 7 on the level of concreteness, estimated age of acquisition (AoA), imageability, concreteness, emotionality and valence, in a web-based behavioral questionnaire. The assessment was done by thirteen naïve respondents that did not partake in any other part of the present study. The concrete words were judged as very concrete [mean rating: 6.5 (sd 0.5)]. The abstract category contained 30 highly-abstract words [mean concreteness: 2.0 (sd 0.9), mean imageability: 2.3 (sd 1.0)] and 29 medium-abstract words [mean concreteness: 3.9 (sd 0.7); mean imageability: 4.1 (sd 0.8)]. Previous studies that highly imaginable word tend to be acquired earlier than words with low imageability [33]. Also in the present stimulus set the estimated AoA for concrete words [mean rating 1.2 (sd 0.3)] was significantly lower [*t* (58) = −9.2, *p* < 0.001] as compared to abstract words [mean rating 2.1 (sd 0.6)]. There was no difference in valence between the word categories [*t* (58) = 1.20, *p* = 9.23].

The concrete words were sub-grouped according to categories that have been derived from specific impairments following brain damage [34–36], namely Animal (e.g., dog, bear), Body part (e.g., hand, foot), Building (e.g., brigde, hospital), Human character (e.g., child, princess), Nature (e.g., island, fire), and Object (e.g., hammer, ball). Each category contained 10 items, with the exception of the Human character category that only contained 9 items. The full list of the stimuli is reported in Supplementary Table S1.

### Corpus-derived Statistical model of semantics

The corpus-derived Statistical model was created using a continuous skip-gram Word2vec-algorithm [16, 17] which looks for co-occurrences between a particular word and the neighboring words (i.e., linguistic context) and represents this information as a N-dimensional vector. The model was trained on a corpus based on a large sample of internet sites in Finnish (1.5 billion words) with negative sampling to approximate the conditional log-likelihood of the model [37]. In the resulting vector space, words that share a similar linguistic context are located close to each other. Here, the vector length used was 300, and a context window of 10 words before and after the stimulus word was used to capture the co-occurrences.

In order to ensure that the SOM visualization of the corpus-derived Statistical model set on the 118 stimulus words reflects the general statistical properties of the corpus data we trained an alternative version of the self-organizing map on all the nouns in the corpus with a frequency > 50 but excluded the 118 stimulus words. We then compared the internal structure of this model to that of the 118-word Statistical model by comparing the location of the projected stimulus words in the SOM structure (see details of the SOM analysis below). The general abstract–concrete dichotomy was also visible in the visualization based on the much larger vocabulary, but the taxonomic class structure was less salient. The results of this control analysis are depicted and described in further detail in the Supplementary Information.

### SOM analysis

We used a self-organizing map [15, 20] (SOM) to evaluate how the different word categories are clustered in the Statistical model. A SOM is an unsupervised artificial neural network, which produces a two dimensional discretized representation of the data; here the respective semantic feature sets. The topography for the SOM was a toroidal grid with 72 nodes (linearly initialized) that is best visualized as a two-dimensional rectangular lattice (9 × 8) where the nodes on opposite edges are connected. With these parameters, the topographic error (0.02) and quantization error (2.2) were small, meaning that the map adequately captures the continuity of the input space with sufficient resolution.

### Experimental design

During the MEG recording the stimulus words were presented in a black mono spaced font (Courier New) on a grey background. Each word was presented for 150 ms followed by a blank screen for 950 ms Between trials, a fixation cross was presented for 1000 ms Each word was presented a total of 20 times, over the course of two one-hour long MEG sessions that took place on separate days. The sessions included breaks of a few minutes every 20 minutes. The order of the stimulus words was randomly determined for each day, so that each stimulus was repeated 10 times each day but words were never repeated back to back.

In order ensure the compliance of the participants, 10 % of the trials were followed by a catch trial, during which the end part of a sentence was presented on the screen and the subject was instructed to determine if the preceding word would make sense as the first word of this sentence. For example, the word ‘beauty’ might be followed by the phrase ‘… is in the eyes of the beholder’ in which case the correct answer would be ‘yes’ as the phrase ‘beauty is in the eyes of the beholder’ is a reasonable sentence.

### MEG and MR measurements

MEG was measured using a whole-head Vectorview MEG device (Elekta Oy, Helsinki, Finland) with 102 triplet sensor elements, each containing two planar gradiometers and one magnetometer. The data was filtered at 0 Hz to 3 Hz sampled at 1000 Hz. Eye movements and blinks were recorded using an electro-oculogram (EOG), configured as pairs of electrodes placed vertically and horizontally around the eyes. The head position with respect to the scanner was determined by four indicator coils placed on the forehead and behind the ears. The head position was measured at the beginning of each 20 minute segment of the recording session. The position of the coils, as well as approx. 60 additional points along the surface of the head, were determined in a coordinate system spanned by three anatomical landmarks (the left and right preauricular points and the nasion) using a 3D Polhemus digitizer (Polhemus, Colchester, VT). The MEG data was co-registered to the anatomical MR images based on the anatomical landmarks and the additional data points.

Anatomical MR images were scanned on a separate day using a 3T MAGNETOM Skyra scanner (Siemens Healthcare, Erlangen, Germany), a standard 20-channel head-neck coil and a T1-weighted MP-RAGE sequence.

### MEG data analysis

The MEG data were preprocessed by aligning head positions from the different data segments and different days into one head position and removing external noise sources using the spatiotemporal signal space separation method [38] in the Elekta Maxfilter software package. Artefactual signals due to eye blinks were suppressed using a PCA approach where 1–2 main components of the average MEG response to blinks were removed from the raw data.

Event-related epochs were extracted from the gradiometer data from 200 ms before to 1000 ms after each word onset and averaged across the multiple presentations of the same item. The event-related responses were baseline-corrected to the interval from −200 ms until the word onset and low-pass filtered at 25 Hz. Any trials where the signal exceeded 3000 fT/cm were removed (max. 1 trial per word).

Source-level estimates were computed using Minimum Norm Estimates (MNE) [39–41] constrained to the cortical surface. The volume conduction model was based on the individual structural MRIs using the Freesurfer software package [42, 43] and modeled as a single-compartment boundary element model with an icosahedron mesh of 2562 vertices in each hemisphere for each participant.

In the inverse solution, currents tangential to the cortical surface were favored by setting the loose orientation constraint parameter to 0.3, and depth-weighting was used to reduce the bias towards superficial sources [44]. The source estimate regularization parameter lambda was set to 0.1. A noise-covariance matrix based on the baseline period across all stimuli was used for noise normalizing of the source estimates, resulting in dynamical statistical parametric maps (dSPMs) [44]. Lastly, the individual source estimates were morphed onto FreeSurfer’s average template brain.

### Zero-shot decoding

In order to determine whether the Statistical model of the semantic space is a good description of the neural responses during word reading, we used a zero-shot decoding machine learning algorithm [21], which was evaluated using a leave-two-out discrimination task.

The zero-shot algorithm was used to learn a linear mapping between the sensor-level MEG evoked responses and the Statistical model. To reduce the dimensionality of the input data, the MEG responses were downsampled by creating 20-ms bins within the time window 0–800 ms relative to the onset of the stimulus presentation, resulting in 40 bins. For each of the 118 stimulus words, the averaged signals for each bin at each of the 204 sensor locations (only the gradiometers were used for decoding) were concatenated into a single vector, yielding a (118 × 8160) input matrix. Ridge regression was used to create a linear mapping between the input matrix and target matrix, i.e., the Statistical model (118 × 300) [45]. The columns of both the input and target matrices were z-transformed before being entered into the linear regression.

The resulting mapping was evaluated by attempting to match two previously unseen segments of MEG data with two unseen stimulus words. To do this, the zero-shot approach employs two steps. First, the algorithm uses the learned mapping between the MEG-data and the individual features to translate the two MEG segments into two predicted feature vectors. The identity of the two unseen stimulus words is then determined by comparing the cosine distance between the predicted vectors and the original Statistical model vectors for these items [22]. This binary discrimination task is carried out for all possible pairs of two stimulus words, using the remaining 116 words for training. For each participant, we report the mean accuracy across all word pairs, which ranges between 50 % (algorithm fails to distinguish between words) and 100 % (successful discrimination between all stimulus words).

To test whether the obtained accuracy scores were significantly higher than chance level, the zero-shot classification procedure was repeated 1000 times on randomly permuted data. Random data was produced by choosing the data of one subject at random and randomizing the assignment between the word labels and the MEG data segments. As p-value, we report the percentage of accuracy scores for the random permutations that equaled or exceeded the accuracy score obtained on real data.

### RSA analysis

Representational similarity analysis (RSA) [23] was performed between the source localized MEG data and Statistical model. For the Statistical model, a single word-to-word dissimilarity matrix (DSM) was created by computing the Pearson correlation *r* across the feature vectors for each possible word pair, and using (1 − *r*) as the dissimilarity score. The values along the diagonal (the dissimilarity between a word and itself) were set to zero.

The MEG data underwent the same downsampling and z-transformation procedure used for the zero-shot learning. Then, for each subject, time bin and source-level vertex, a word-to-word DSM was formed using a searchlight approach: the signal at all vertices within a 3 cm radius of the vertex under consideration were assembled into a vector. These vectors were then compared for all possible combinations of two words using Pearson correlation, with (1 − *r*) as dissimilarity score.

The RSA maps for each subject and each feature set were obtained by comparing the MEG-based DSMs with the feature-set DSMs using Spearman rank correlation. Finally, the RSA maps were analyzed across subjects using a cluster permutation test [24] with a cluster threshold of *p* = 0.01 and the significance threshold for the permuted randomly shuffled data distribution set to *p* = 0.05. The number of permutations used to create the random distribution of the data was 5000. Any clusters with a corresponding cluster-*t*-value that was lower than 95 % of the randomly obtained cluster-*t*-values were pruned from the RSA maps. The remaining clusters were deemed significant (*p* ≤ 0.05).

To aid the interpretation of the main RSA, an additional RSA was calculated between the MEG data and a separate model quantifying only the abstract – concrete dimension (Abstractness model; see below). This additional RSA was computed in the same manner as the main RSA between the MEG data and the Statistical model, with the exception that the Euclidean distance was used as the distance metric in the word-to-word DSM of the one-dimensional Abstractness model.

### Abstractness model

In the RSA analysis, the main results were interpreted in light of a complementary RSA between the MEG data and an Abstractness model capturing the abstract–concrete dimension of the stimulus words (Abstractness model RSA). The Abstractness model was derived from a behavioral web-based questionnaire answered by 10 naïve respondents (that did not take part of the stimulus assessment questionnaire). Each respondent judged the 118 stimulus words with respect to six different taxonomic categories (Animal, Body part, Building, Human character, Nature, Object) and abstractness on a scale from 1–7 (1 = does not belong to this category, 7 = a typical example to this category). From this data set, we extracted the abstractness scale to be used as an Abstractness model.

### Data and code availability

The text corpus containing 1.5 billion Finnish words used to derive the statistical model cannot be publicly distributed due to the Finnish copyright law limitations. It is available upon request for research purposed, for contact information see http://bionlp.utu.fi/finnish-internet-parsebank.html. The word2vec models used in this study (derived from the abovementioned corpus), together with the custom code used in the study can be accessed from https://version.aalto.fi/gitlab/BrainDecode/zeroshotdecoding. The stimulus words are publicly available and listed in the supplementary information. The MEG and MRI data are available upon reasonable requests from the authors; the data is not publicly available as it contains information that could compromise the participant privacy and consent.

## Acknowledgements

We would like to express our gratitude to Jenna Kanerva and Filip Ginter at the University of Turku for development of the Finnish language Word2vec model as well as to Enrico Glerean and Gus Sudre for sharing code used in the study. This research was funded by the Academy of Finland (grant #287474 to A.H., #286070 to S. L. K., #310988 to MvV, and #255349, #256459 and #283071 to R.S.), the Aalto Brain Center (MvV and T.L-K.) as well as the Sigrid Jusélius Foundation (R.S.). Computational resources were provided by the Aalto Science-IT project and the CSC - IT center for science Ltd..

## Supplementary Information

### Breakdown of decoding prediction

In order to understand if any underlying systematics in the stimulus selection was driving the performance of the zero-shot decoding model, we looked at the pairwise comparisons in the leave-two-out cross validation scheme. Figure S1 shows the performance of each combination of item-pairs averaged across all participants. As no clear categorical pattern emerges, we conclude that the prediction accuracy is not driven by a specific category of the stimulus words.

**Figure S1:**
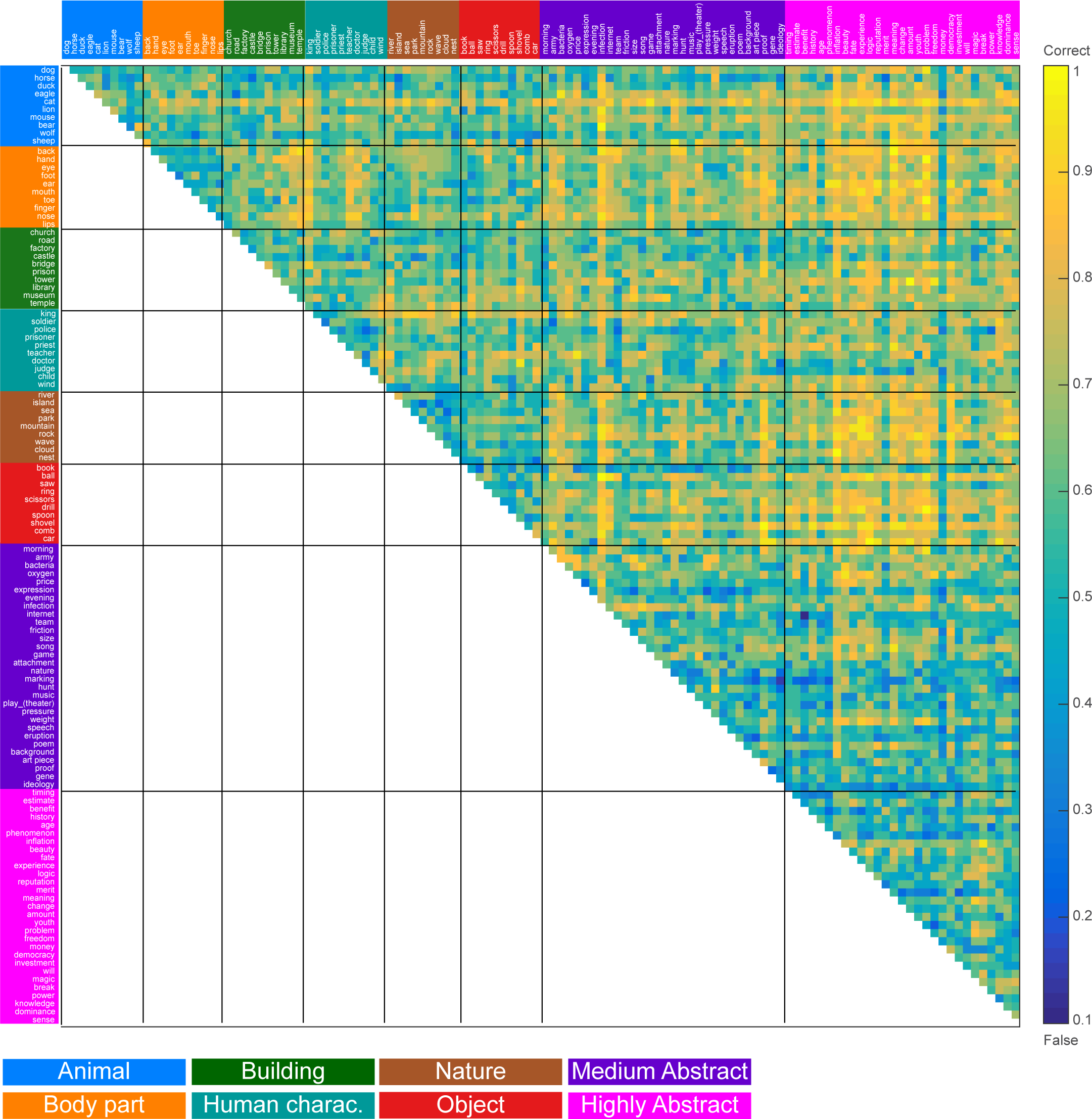
Prediction accuracy for each stimulus-item pair, averaged across participants.

### SOM control analysis

In order to ensure that the SOM visualization of the corpus-derived Statistical model set (on 118 words) reflects the general statistical properties of the corpus data, and is not dependent on the particular set of stimulus words used here, we trained a supplementary SOM on a Corpus-wide statistical model. This model included all words with a frequency > 50 (N=319523 words), but omitted the 118 stimulus words used in the MEG study. The SOM was trained using the same parameters as the main Statistical model SOM, i.e., on toroidal grid with 72 nodes. For the Corpus-wide statistical model SOM the topographic error was 0.08 and the quantization error 2.8. The map was evaluated by visualizing the location of the stimulus word in the model.

Similarly to the main Statistical model SOM (see Figure 1 in the main article) the abstract and concrete words largely formed distinct groups also in the Corpus-wide statistical model SOM (Figure S2). Especially words classified as ‘Body part’ were clearly distinct from the other word categories. Two nodes in the center of the map contained words from several taxonomic categories and included both abstract and concrete words. These nodes seemed to be based on thematic relationships between the words.

**Figure S2:**
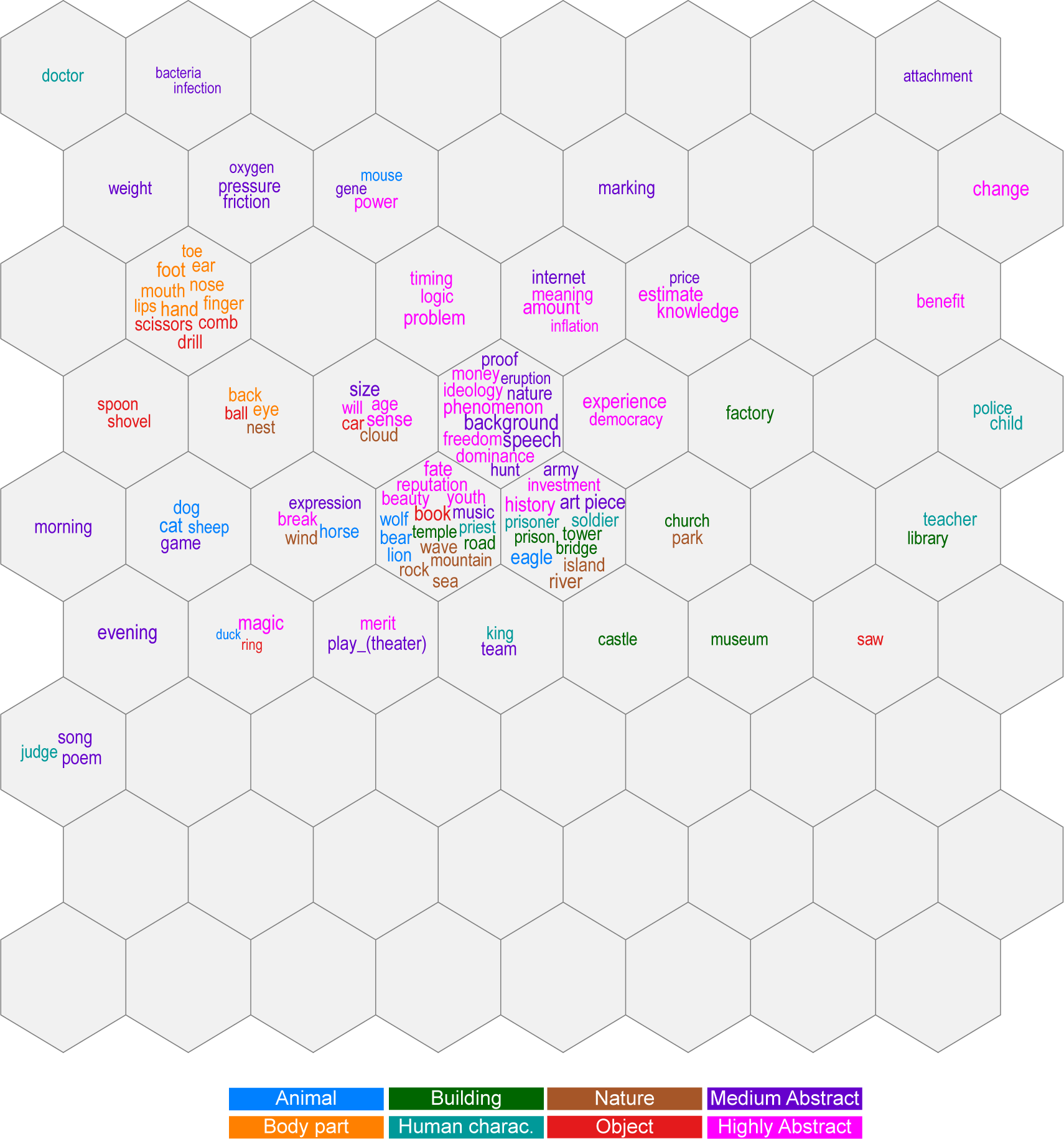
Visualization of the stimulus words in a SOM trained on a Corpus-wide statistical model. The size of the each word denotes the distance to the node center (the larger the font, the closer to the node center the word lies) and the color the predefined category class of the stimulus words

**Table S1. Supplementary Table 1.** Full list of stimuli

| STIMULUS | TRANSLATION | CATEGORY | STIMULUS | TRANSLATION | CATEGORY |
| --- | --- | --- | --- | --- | --- |
| koira | dog | Animal | aamu | morning | Medium Abstract |
| hevonen | horse | Animal | armeija | army | Medium Abstract |
| ankka | duck | Animal | bakteeri | bacteria | Medium Abstract |
| kotka | eagle | Animal | happi | oxygen | Medium Abstract |
| kissa | cat | Animal | hinta | price | Medium Abstract |
| leijona | lion | Animal | ilme | expression | Medium Abstract |
| hiiri | mouse | Animal | ilta | evening | Medium Abstract |
| karhu | bear | Animal | infektio | infection | Medium Abstract |
| susi | wolf | Animal | internet | internet | Medium Abstract |
| lammas | sheep | Animal | joukkue | team | Medium Abstract |
| selkä | back | Body part | kitka | friction | Medium Abstract |
| käsi | hand | Body part | koko | size | Medium Abstract |
| silmä | eye | Body part | laulu | song | Medium Abstract |
| jalka | foot | Body part | leikki | game | Medium Abstract |
| korva | ear | Body part | liite | attachment | Medium Abstract |
| suu | mouth | Body part | luonto | nature | Medium Abstract |
| varvas | toe | Body part | merkintä | marking | Medium Abstract |
| sormi | finger | Body part | metsästys | hunt | Medium Abstract |
| nenä | nose | Body part | musiikki | music | Medium Abstract |
| huulet | lips | Body part | näytelmä | play (theater) | Medium Abstract |
| kirkko | church | Building | paine | pressure | Medium Abstract |
| tie | road | Building | paino | weight | Medium Abstract |
| tehdas | factory | Building | puhe | speech | Medium Abstract |
| linna | castle | Building | purkaus | eruption | Medium Abstract |
| silta | bridge | Building | runo | poem | Medium Abstract |
| vankila | prison | Building | tausta | background | Medium Abstract |
| torni | tower | Building | teos | artwork | Medium Abstract |
| kirjasto | library | Building | todiste | proof | Medium Abstract |
| museo | museum | Building | geeni | gene | Medium Abstract |
| temppeli | temple | Building | aate | ideology | Highly Abstract |
| kuningas | king | Human character | ajoitus | timing | Highly Abstract |
| sotilas | soldier | Human character | arvio | estimate | Highly Abstract |
| poliisi | police | Human character | etu | benefit | Highly Abstract |
| vanki | prisoner | Human character | historia | history | Highly Abstract |
| pappi | priest | Human character | ikä | age | Highly Abstract |
| opettaja | teacher | Human character | ilmiö | phenomenon | Highly Abstract |
| lääkäri | doctor | Human character | inflaatio | inflation | Highly Abstract |
| tuomari | judge | Human character | kauneus | beauty | Highly Abstract |
| lapsi | child | Human character | kohtalo | fate | Highly Abstract |
| tuli | wind | Nature | kokemus | experience | Highly Abstract |
| joki | river | Nature | logiikka | logic | Highly Abstract |
| saari | island | Nature | maine | reputation | Highly Abstract |
| meri | sea | Nature | meriitti | merit | Highly Abstract |
| puisto | park | Nature | merkitys | meaning | Highly Abstract |
| vuori | mountain | Nature | muutos | change | Highly Abstract |
| kallio | rock | Nature | määrä | amount | Highly Abstract |
| aalto | wave | Nature | nuoruus | youth | Highly Abstract |
| pilvi | cloud | Nature | ongelma | problem | Highly Abstract |
| pesä | nest | Nature | vapaus | freedom | Highly Abstract |
| kirja | book | Object | raha | money | Highly Abstract |
| pallo | ball | Object | demokratia | democracy | Highly Abstract |
| saha | saw | Object | sijoitus | investment | Highly Abstract |
| sormus | ring | Object | tahto | will | Highly Abstract |
| sakset | scissors | Object | taika | magic | Highly Abstract |
| pora | drill | Object | tauko | break | Highly Abstract |
| lusikka | spoon | Object | teho | power | Highly Abstract |
| lapio | shovel | Object | tieto | knowledge | Highly Abstract |
| kampa | comb | Object | valta | dominance | Highly Abstract |
| auto | car | Object | järki | sense | Highly Abstract |

## References

[1] J. R. Binder, L. L. Conant, C. J. Humphries, L. Fernandino, S. B. Simons, M. Aguilar, and R. H. Desai. “Toward a brain-based componential semantic representation.” In: Cognitive Neuropsychology (2016), pp. 1–45. doi: 10.1080/02643294.2016.1147426.

[2] M. Kiefer and F. Pulvermüller. “Conceptual representations in mind and brain: theoretical developments, current evidence and future directions.” In: Cortex 48.7 (2012), pp. 805–25. doi: 10.1016/j.cortex.2011.04.006.

[3] A. Martin. “The representation of object concepts in the brain.” In: Annu. Rev. Psychol. 58 (2007), pp. 25–45. doi: 10.1146/annurev. psych.57.102904.190143.

[4] G. Vigliocco and D. P. Vinson. “Semantic Representation.” In: The ‘Oxford handbook of psycholinguistics, ed. by M. G. Gaskell. Oxford University Press, 2009, pp. 195–217.

[5] G. Hollis and C. Westbury. “The principals of meaning: Extracting semantic dimensions from co-occurrence models of semantics.” In: Psychon. Bull. Rev. 23.6 (2016), pp. 1744–1756. doi: 10. 3758/s13423-016-1053-2.

[6] Donald O. Hebb. The organization of behavior. New York: Wiley, 1949.

[7] K. Friston. “The history of the future of the Bayesian brain.” In: Neuroimage 62.2 (2012), pp. 1230–3. doi: 10.1016/j.neuroimage.2011.10.004.

[8] George Zipf. Human Behavior and the Principle of Least Effort: An Introduction to Human Ecology. Addison-Wesley Press, 1949.

[9] Carlton T. James. “The role of semantic information in lexical decisions.” In: J. Exp. Psychol. Hum. Percept. Perform. 1.2 (1975), p. 130.

[10] Jamie Reilly, Jonathan E. Peelle, and Murray Grossman. “A unitary semantics account of reverse concreteness effects in semantic dementia.” In: Brain. Lang. 103.1 (2007), pp. 86–87.

[11] E. K. Warrington. “The selective impairment of semantic memory.” In: Q J Exp Psychol 27.4 (1975), pp. 635–57. doi: 10.1080/14640747508400525.

[12] J. Wang, J. A. Conder, D. N. Blitzer, and S. V. Shinkareva. “Neural representation of abstract and concrete concepts: a meta-analysis of neuroimaging studies.” In: Hum. Brain Mapp. 31.10 (2010), pp. 1459–68. doi: 10.1002/hbm.20950.

[13] Phillip J. Holcomb, John Kounios, Jane E. Anderson, and W. Caroline West. “Dual-coding, context-availability, and concreteness effects in sentence comprehension: An electrophysiological investigation.” In: J. Exp. Psychol. Learn. Mem. Cogn. 25.3 (1999), pp. 721–742.

[14] Hsu-Wen Huang, Chia-Lin Lee, and Kara D. Federmeier. “Imagine that! ERPs provide evidence for distinct hemispheric contributions to the processing of concrete and abstract concepts.” In: Neuroimage 49.1 (2010), pp. 1116–1123.

[15] P. Teuvo Kohonen. “Self-organized formation of topologically correct feature maps.” In: Biol. Cybern. 43.1 (1982), pp. 59–69.

[16] Tomas Mikolov, Wen-tau Yih, and Geoffrey Zweig. “Linguistic regularities in continuous space word representations.” In: Human Language Technologies. Proceedings of the 2013 conference of the North American chapter of the association for computational linguistics. 2013, pp. 746–751.

[17] Tomas Mikolov, Kai Chen, Greg Corrado, and Jeffrey Dean. “Efficient estimation of word representations in vector space.” In: arXiv preprint arXiv:1301.3781 (2013).

[18] Simon De Deyne, Steven Verheyen, and Gert Storms. “Structure and organization of the mental lexicon: A network approach derived from syntactic dependency relations and word associations.” In: Towards a theoretical framework for analyzing complex linguistic networks. Springer, 2016, pp. 47–79.

[19] Emilie L. Lin and Gregory L. Murphy. “Thematic relations in adults’ concepts.” In: J. Exp. Psychol. Gen. 130.1 (2001), p. 3.

[20] Timo Honkela, Ville Pulkki, and Teuvo Kohonen. “Contextual relations of words in Grimm tales analyzed by self-organizing map.” In: Proceedings of the International Conference on Artificial Neural Networks (ICANN-95). Vol. 2. EC2 et Cie Paris, 1995, pp. 3–7.

[21] Mark Palatucci, Dean Pomerleau, Geoffrey E. Hinton, and Tom M. Mitchell. “Zero-shot learning with semantic output codes.” In: Advances in neural information processing systems. 2009, pp. 1410–1418.

[22] G. Sudre, D. Pomerleau, M. Palatucci, L. Wehbe, A. Fyshe, R. Salmelin, and T. Mitchell. “Tracking neural coding of perceptual and semantic features of concrete nouns.” In: Neuroimage 62.1 (2012), pp. 451–63. doi: 10.1016/j.neuroimage.2012.04.048.

[23] Nikolaus Kriegeskorte, Marieke Mur, and Peter A. Bandettini. “Representational similarity analysis-connecting the branches of systems neuroscience.” In: Front. Syst. Neurosci. 2 (2008), p. 4.

[24] E. Maris and R. Oostenveld. “Nonparametric statistical testing of EEG-and MEG-data.” In: J. Neurosci. Methods 164.1 (2007), pp. 177–90. doi: 10.1016/j.jneumeth.2007.03.024.

[25] Jing Wang, Laura B. Baucom, and Svetlana V. Shinkareva. “Decoding abstract and concrete concept representations based on singlein-validtrial fMRI data.” In: Hum. Brain Mapp. 34.5 (2013), pp. 1133–1147.

[26] A. Clarke and L. K. Tyler. “Object-Specific Semantic Coding in Human Perirhinal Cortex.” English. In: J. Neurosci. 34.14 (2014), pp. 4766–4775. doi: 10.1523/Jneurosci.2828-13.2014.

[27] Jeffrey R. Binder, Chris F. Westbury, Kristen A. McKiernan, Edward T. Possing, and David A. Medler. “Distinct brain systems for processing concrete and abstract concepts.” In: J. Cogn. Neurosci. 17.6 (2005), pp. 905–917.

[28] K. Patterson, P. J. Nestor, and T. T. Rogers. “Where do you know what you know? The representation of semantic knowledge in the human brain.” English. In: Nat. Rev. Neurosci. 8.12 (2007), pp. 976–987. doi: 10.1038/nrn2277.

[29] C. J. Price. “A review and synthesis of the first 20 years of PET and fMRI studies of heard speech, spoken language and reading.” In: Neuroimage 62.2 (2012), pp. 816–47. doi: 10.1016/j.neuroimage.2012.04.062.

[30] F. Pulvermuller, F. Kherif, O. Hauk, B. Mohr, and I. Nimmo-Smith. “Distributed cell assemblies for general lexical and category-specific semantic processing as revealed by fMRI cluster analysis.” In: Hum. Brain Mapp. 30.12 (2009), pp. 3837–50. doi: 10.1002/hbm.20811.

[31] Leonardo Fernandino, Colin J. Humphries, Mark S. Seidenberg, William L. Gross, Lisa L. Conant, and Jeffrey R. Binder. “Predicting brain activation patterns associated with individual lexical concepts based on five sensory-motor attributes.” In: Neuropsychologia 76 (2015), pp. 17–26.

[32] I. Simanova, P. Hagoort, R. Oostenveld, and M. A. van Gerven. “Modality-independent decoding of semantic information from the human brain.” In: Cereb. Cortex 24.2 (2014), pp. 426–34. doi: 10. 1093/cercor/bhs324.

[33] H. Stadthagen-Gonzalez and C. J. Davis. “The Bristol norms for age of acquisition, imageability, and familiarity.” In: Behav Res Methods 38.4 (2006), pp. 598–605.

[34] Elizabeth K. Warrington and Tim Shallice. “Category specific semantic impairments.” In: Brain 107.3 (1984), pp. 829–853.

[35] Alfonso Caramazza and Jennifer R. Shelton. “Domain-specific knowledge systems in the brain: The animate-inanimate distinction.” In: J. Cogn. Neurosci. 10.1 (1998), pp. 1–34.

[36] Giuseppe Sartori, Michele Miozzo, and Remo Job. “Category-specific naming impairments? Yes.” In: The Quarterly Journal of Experimental Psychology Section A 46.3 (1993), pp. 489–504.

[37] Jenna Kanerva, Juhani Luotolahti, Veronika Laippala, and Filip Ginter. “Syntactic N-gram collection from a large-scale corpus of internet Finnish.” In: Frontiers in Artificial Intelligence and Applications. Vol. 268. IOS Press, 2014, pp. 184–191. isbn: 1-61499-442-0. doi: 10.3233/978-1-61499-442-8-184.

[38] S. Taulu and J. Simola. “Spatiotemporal signal space separation method for rejecting nearby interference in MEG measurements.” In: Phys. Med. Biol. 51.7 (2006), pp. 1759–68. doi: 10.1088/0031-9155/51/7/008.

[39] A. Gramfort, M. Luessi, E. Larson, D. A. Engemann, D. Strohmeier, C. Brodbeck, R. Goj, M. Jas, T. Brooks, L. Parkkonen, and M. Hamalainen. “MEG and EEG data analysis with MNE-Python.” In: Front. Neurosci. 7 (2013), p. 267. doi: 10.3389/fnins.2013.00267.

[40] A. Gramfort, M. Luessi, E. Larson, D. A. Engemann, D. Strohmeier, C. Brodbeck, L. Parkkonen, and M. S. Hamalainen. “MNE software for processing MEG and EEG data.” In: Neuroimage 86 (2014), pp. 446–60. doi: 10.1016/j.neuroimage.2013.10.027.

[41] M. S. Hämäläinen and R. J. Ilmoniemi. “Interpreting magnetic fields of the brain: minimum norm estimates.” In: Med. Biol. Eng. Comput. 32.1 (1994), pp. 35–42.

[42] A. M. Dale, B. Fischl, and M. I. Sereno. “Cortical surface-based analysis. I. Segmentation and surface reconstruction.” In: Neuroimage 9.2 (1999), pp. 179–94. doi: 10.1006/nimg.1998.0395.

[43] B. Fischl, A. Liu, and A. M. Dale. “Automated manifold surgery: constructing geometrically accurate and topologically correct models of the human cerebral cortex.” In: IEEE Trans. Med. Imaging. 20.1 (2001), pp. 70–80. doi: 10.1109/42.906426.

[44] Anders M. Dale, Arthur K. Liu, Bruce R. Fischl, Randy L. Buckner, John W. Belliveau, Jeffrey D. Lewine, and Eric Halgren. “Dynamic statistical parametric mapping: combining fMRI and MEG for high-resolution imaging of cortical activity.” In: Neuron 26.1 (2000), pp. 55–67.

[45] Fabian Pedregosa, Gaël Varoquaux, Alexandre Gramfort, Vincent Michel, Bertrand Thirion, Olivier Grisel, Mathieu Blondel, Peter Prettenhofer, Ron Weiss, and Vincent Dubourg. “Scikit-learn: Machine learning in Python.” In: Journal of machine learning research 12.Oct (2011), pp. 2825–2830.

